# Pudendal, but not tibial, nerve stimulation modulates vulvar blood perfusion in anesthetized rodents

**DOI:** 10.1101/2022.03.05.483101

**Authors:** Elizabeth C. Bottorff, Tim M. Bruns

**Affiliations:** University of Michigan, Biomedical Engineering Department, Ann Arbor, MI, USA; University of Michigan, Biointerfaces Institute, Ann Arbor, MI, USA

**Keywords:** blood flow, laser speckle imaging, neuromodulation, pudendal nerve, tibial nerve, vulva

## Abstract

**Introduction and Hypothesis:** Preclinical studies have shown that neuromodulation can increase vaginal blood perfusion, but the effect on vulvar blood perfusion is unknown. We hypothesized that pudendal and tibial nerve stimulation could evoke an increase in vulvar blood perfusion.

**Methods:** We used female Sprague-Dawley rats for non-survival procedures under urethane anesthesia. We measured perineal blood perfusion in response to twenty-minute periods of pudendal and tibial nerve stimulation using laser speckle contrast imaging (LSCI). After a thoracic-level spinalization and a rest period, we repeated each stimulation trial. We calculated average blood perfusion before, during, and after stimulation for three perineal regions (vulva, anus, and inner thigh), for each nerve target and spinal cord condition.

**Results:** We observed a significant increase in vulvar, anal, and inner thigh blood perfusion during pudendal nerve stimulation in spinally intact and spinalized rats. Tibial nerve stimulation had no effect on perineal blood perfusion for both spinally intact and spinalized rats.

**Conclusions:** This is the first study to examine vulvar hemodynamics with LSCI in response to nerve stimulation. This study demonstrates that pudendal nerve stimulation modulates vulvar blood perfusion, indicating the potential of pudendal neuromodulation to improve genital blood flow as a treatment for women with sexual dysfunction. This study provides further support for neuromodulation as a treatment for women with sexual arousal disorders. Studies in unanesthetized animal models with genital arousal disorders are needed to obtain further insights into the mechanisms of neural control over genital hemodynamics.

**Brief Summary:** In an anesthetized rodent model, electrical stimulation of the pudendal nerve will drive increases in vulvar blood perfusion while tibial nerve stimulation will not.

## Introduction

Female sexual health is an important determinant in quality of life, contributing to both increased meaning in life and general well-being [1]. Unfortunately, approximately 40% of women suffer from female sexual dysfunction (FSD) [2]. Female sexual dysfunction can present in multiple domains, which can be thought of as physiological (arousal, lubrication, orgasm, pain) and psychological (satisfaction and desire). Due to limited research regarding the etiology of FSD and basic female anatomy, current treatment options for FSD are very limited, especially for women who have deficits in the physiological domains such as arousal [3]. Bremelanotide and flibanserin, both approved to treat hypoactive sexual desire disorder (HSDD), have shown mild efficacy in improving sexual desire and arousal [4, 5]. However, bremelanotide has been associated with a high incidence of adverse events [6] and a review of flibanserin, suggests that it has minimal clinical benefit [7]. Sildenafil citrate has been studied as a treatment for female arousal disorders. One study of sildenafil demonstrated an increase in clitoral vasocongestion and an association with increased sexual satisfaction [8]. Sildenafil was also used to study vaginal vasocongestion in treatment and placebo groups during erotic visual stimulation [9]. Although this study reported significant increases in vasocongestion for the treatment group, there were no differences in subjective arousal between the sildenafil and placebo groups. The lack of consistent improvements in subjective arousal as well as a high incidence of adverse events led to sildenafil citrate no longer being pursued as a treatment option [10].

Neuromodulation is a potential treatment option for women with FSD. Sacral neuromodulation (SNM) is a standard treatment for overactive bladder and fecal incontinence [11, 12]. SNM delivers electrical stimulation to sacral nerves, which contain somatic and sympathetic nerve fibers of the pelvic organs. Clinical studies have found that sexual function can improve in women receiving SNM for bladder function [13, 14]. Similar improvements in sexual function have been seen in a clinical study that used tibial nerve stimulation to treat bladder dysfunction [15]. The sexual health benefits of neuromodulation have been studied in women without concomitant pelvic disorders. In a recent study, transcutaneous electrical nerve stimulation (TENS) of the tibial nerve or dorsal genital nerve (DGN), a distal branch of the pudendal nerve, improved survey-reported female sexual function index (FSFI) scores, a standardized metric for evaluating female sexual function [16]. These subjects reported an improvement in their overall FSFI score and their individual arousal and orgasm sub-scores. Subjects receiving DGN stimulation also saw improvements in sexual satisfaction. There is a lack of physiological measurements of sexual health and so the mechanisms by which pudendal and tibial nerve stimulation improve sexual function are not completely understood. One possible mechanism is that nerve stimulation modulates genital blood flow, which is necessary to facilitate vasocongestion, lubrication, and sexual receptivity [17]. This is supported by preclinical studies showing that stimulation of the tibial nerve for 30 minutes can result in a transient increase in vaginal blood flow 20-35 minutes after stimulation onset [18]. Preclinical studies have also shown an increase in vaginal blood flow in response to pudendal nerve stimulation, with peak responses occurring anywhere from 1-2 minutes [19] or up to 30 minutes [20] after stimulation onset depending on stimulation parameters. Additionally, it is well established that stimulation of the somatic pudendal nerve and tibial nerve can modulate spinal control over the bladder and bowel, leading to improvement in dysfunctional states [21]. We hypothesized that a similar spinal reflex exists for genital vasocongestion. Both pudendal and tibial nerve stimulation have the potential to improve sexual function and have specific translational advantages, including the ease of access and existing clinical use for pelvic dysfunctions [15, 22], however a greater understanding of the underlying mechanisms and neural circuitry is needed.

Female sexual dysfunction often presents in women with spinal cord injuries (SCI). Women with SCI retain varying aspects of sexual function, depending on their injury type. For example, research has shown that sensory impairment to the T11-L2 dermatomes, but not injury level, is associated with decreased genital arousal [23]. Although paraplegic patients report that regaining sexual function after SCI is a priority [24], women with SCI have the same FSD treatment options as non-neurogenic women. These treatment options do not consider the impact of neurological dysfunction. Furthermore, neuromodulation via tibial nerve stimulation may need supraspinal pathways [21] which would limit utility for SCI women depending on their injury severity. Once the neural pathways controlling genital arousal are better understood, clinicians can use knowledge of the patient’s injury to determine if electrical stimulation is able modulate these pathways to improve genital arousal in women with neurogenic FSD. However, it is first necessary to identify how different nerve targets of neuromodulation modulate genital arousal. In healthy women, genital arousal is a physiological response characterized by an increase in genital blood flow, engorgement of the genitals, and the production of lubrication [17]. Most research studying female sexual responses use vaginal photoplethysmography to measure genital arousal [25]. However, some researchers are turning towards more non-invasive methods such as laser doppler imaging or laser speckle contrast imaging (LSCI). These laser-based methodologies are becoming more useful in measuring female sexual arousal because they show a higher concordance between genital and subjective arousal[26]. However, the importance of genital-subjective arousal synchrony is not well understood [3] and some models for the sexual response cycle suggest that genital arousal is necessary for satisfactory sexual intercourse, regardless of subjective arousal [27].

In this preclinical study we used LSCI to measure blood perfusion changes in the perineal region of female rats in response to tibial and pudendal nerve stimulation. Additionally, we looked at how this blood perfusion response changes after thoracic level spinalization. Although LSCI has been used to assess genital arousal in clinical studies [26, 28], this is the first time it has been used to measure perineal blood perfusion in animals. Our main outcome measures for this study were changes in vulvar, anal, and inner thigh blood perfusion from baseline in response to nerve stimulation. We hypothesized that pudendal nerve stimulation would drive larger increases in vulvar blood perfusion than tibial nerve stimulation. We also hypothesized that spinal cord transection would decrease the perineal blood perfusion response to tibial nerve stimulation, but not pudendal nerve stimulation.

## Materials and Methods

### Animal Surgery

All experimental procedures were approved by the University of Michigan Institutional Animal Care and Use Committee (IACUC) in accordance with the National Institutes of Health’s guidelines for the care and use of laboratory animals. We designed the primary experimental protocol prior to starting the study. The study protocol was not pre-registered. We performed non-survival procedures on 17 nulliparous female Sprague-Dawley rats (Charles River Breeding Labs, Wilmington, MA, USA) weighing 0.25 to 0.30 kilograms. Female Sprague-Dawley rats are a common preclinical model for studying sexual arousal because their physiological response to sexual stimuli is similar to that found in humans (e.g., increases in vaginal blood flow, length, and pressure) [29]. Additionally, the neuroanatomical pathways of the pelvic, hypogastric, and pudendal nerves, all which mediate genital arousal and sensation, have been well studied in rats [30]. We performed a power analysis using a Wilcoxon signed-rank test with α = 0.05, power = 0.80, and an effect size estimate from genital perfusion changes in a study with women [31] that resulted in a sample size of 9. We targeted 15 rats to account for the potential for higher variability in animal genital perfusion responses and possible animal deaths under anesthesia. Ultimately we had 10 animals that completed the full set of experiments, which matches prior animal studies with similar objectives [20, 32]. Animals were housed in ventilated cages under controlled temperature, humidity, and photoperiod (12-h light/dark cycle), and provided laboratory chow (5L0D, LabDiet, St. Louis, MO, USA). Animals were anesthetized using intraperitoneal urethane (1.5 g/kg), a commonly used anesthetic in rodent surgeries studying pelvic organ function [19, 33]. Sufficient anesthetic depth was confirmed once the animal no longer had a toe pinch response. We used a heating pad to maintain body temperature and monitored vital signs (heart rate, respiration rate, and oxygen saturation levels) every 15 minutes. We used vital signs as humane endpoints. We performed vaginal cytology prior to surgical access to determine estrous stage, to examine if one stage (e.g. estrus) better facilitates a perineal blood flow response.

With the rat in the prone position, we made a dorsal midline incision through the skin 3-4 cm rostral to the base of the tail. Then we extended the incision laterally on the right side, and the ischiorectal fossa was separated. We used retractors to keep the musculature open and isolated the pudendal nerve using forceps. We placed a bipolar nerve cuff with stranded stainless-steel wire electrodes (400 μm diameter; Cooner Wire Co, Chatsworth, CA, SA) and silicone elastomer tubing (0.5-mm inner diameter; Dow Corning, Midland, MI, USA) around both sensory and motor branches of the pudendal nerve within the ischiorectal fossa, proximal to the division of the motor branch into its dorsal and ventral branches [34]. We then closed the incision and moved the animal to a supine position. We then placed a percutaneous electromyogram (EMG) wire (stainless steel, 50 μm, MicroProbes, Gaithersburg, MD) subcutaneously, parallel to the tibial nerve and ipsilateral to the pudendal nerve cuff. We measured perineal blood perfusion with a laser speckle contrast imaging (LSCI) system (MOORFLPI-2, Moor Instruments, Wilmington, DE). We aimed the laser at the perineal region, angled downward at 25 degrees, and positioned such that the path of the laser traveled 15 to 20 cm to the animal (Figure 1a).

**Figure 1.**
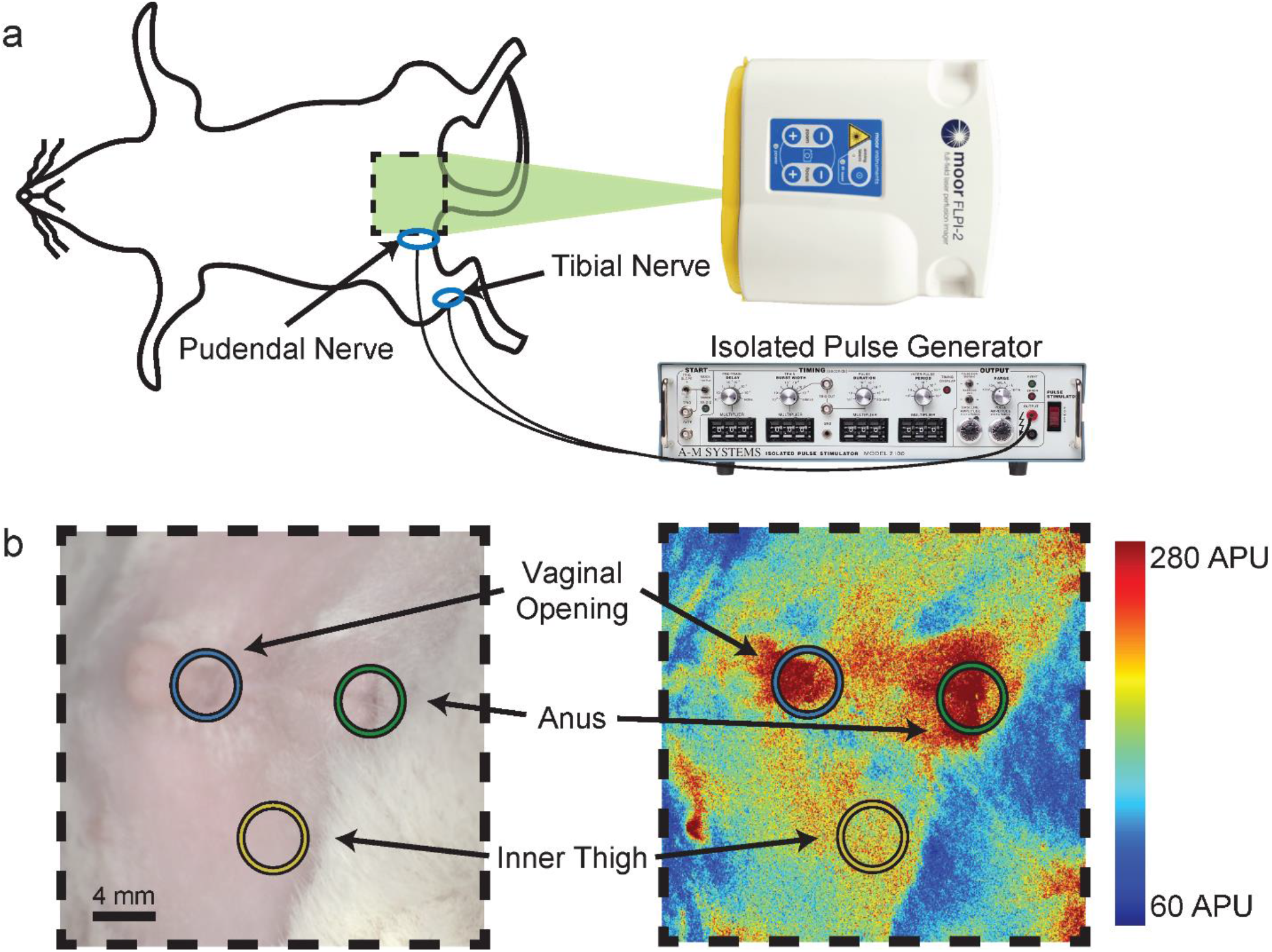
Experimental set-up. Figure 1a shows a diagram of the LSCI with the laser aimed at the vaginal vestibule of a rat lying in the supine position. Figure 1b shows an example of a laser speckle image live (left) and produced as a heatmap (right) with ROIs drawn around the vaginal vestibule, anus, and a region in the inner thigh in blue, green, and yellow respectively.

### Experimental Protocol

We delivered stimulation using an isolated pulse generator (Model 2100, A-M Systems, Sequim, WA). We determined the motor threshold (MT) for stimulating the pudendal and tibial nerves by slowly increasing the stimulation amplitude until a motor response (anal or toe twitch respectively) was observed. Each animal went through a series of 45-minute trials. For each trial, we used LSCI to measure perineal blood perfusion at the maximum sampling rate of 0.25 Hz, using the temporal processing settings of the MOORFLPI-2, for 5 minutes prior to stimulation, during 20 minutes of stimulation, and for 20 minutes after stimulation. We delivered stimulation to one of the nerve targets using biphasic, rectangular pulses (0.2-ms pulse width) at 20 Hz and twice the MT. We repeated the recording protocol for the other nerve target. After both trials, we placed the rat in the prone position and performed a laminectomy. We used a rostral/caudal incision to expose T8-10 level vertebrae. After removal of the dorsal processes, we used a scalpel blade to transect the spinal cord. We closed the musculature with sutures and closed the skin with skin staples. We then moved the rat to the supine position and allowed it to rest for at least 15 minutes. Next, we re-assessed the MT and repeated the trials in the same order as the intact spinal cord trials. After all experimental procedures were complete, we euthanized the animal with an intraperitoneal injection of sodium pentobarbital (250-300 mg/kg). In the first three rats, tibial nerve stimulation was performed first. We alternated the order of the first stimulated nerve across subsequent animals, occasionally using sequential animals with pudendal nerve stimulation first, to arrive at balanced groups upon study termination. Investigators were not blind to stimulation order.

We added two sets of supplemental experiments to the protocol after the study began. In some initial experiments, we saw the blood perfusion signal spontaneously increase over 100 APU for 1 to 3 minutes, at 5-to-6-minute intervals. These increases were too abrupt to be considered a physiological response in blood perfusion, so we performed another set of experiments to examine LSCI data sampling (n = 2). The experimental procedure in these experiments followed the same set of 4 trials (pudendal and tibial stimulation, before and after spinal cord transection), except that the blood perfusion was sampled at 1 Hz or 25 Hz using the spatial processing settings instead of the maximum sampling rate of 0.25 Hz with the temporal processing setting. In the second set of supplemental experiments (n = 3), we transected the pudendal nerve next to the cuff electrode between stimulation trials with an intact spinal cord. We looped a suture string around the nerve during cuff placement and pulled it tightly to cut the nerve. We transected the pudendal nerve a different way in each of these experiments: once proximal to the cuff, once distal to the cuff, and once distal to the cuff with a partial and then complete distal cut.

### Data Analysis

We performed all data and statistical analyses using moorFLPI-2 Research Software (Software-MOORFLPI2-3VX, Moor Instruments, Wilmington, DE) and MATLAB (Mathworks, Natick, MA, USA). We drew regions of interest (ROIs) around the vulva, the anus, and a region on the inner thigh for each LSCI trial analysis in moorFLPI-2. We drew a circular ROI centered on the vaginal orifice, with a diameter equal to the width of the tissue mound surrounding the urethra. We then centered two identical sized ROIs on the anus and the inner thigh on the contralateral side of where stimulation was delivered, such that the three ROIs formed an equilateral triangle (Figure 1b). We calculated average perfusion units within each ROI (vulvar, anal, and inner thigh) and extracted the signal for each 45-minute trial. Each signal is referred to as vulvar blood perfusion (VBP), anal blood perfusion (ABP), and inner thigh blood perfusion (ITBP). We observed an artifact when stimulation was turned on and removed it by eliminating the data from 20 seconds before to 60 seconds after stimulation onset. We then calculated the temporal mean for each ROI before, during, and after stimulation. We used a linear regression to determine the impact of estrous stage, stimulation order, experiment number, and weight on the results. Estrous stage and stimulation order were coded as binary values in this analysis. We made comparisons between average VBP, ABP, and ITBP before and after spinalization as well as across stimulation epochs (before, during, and after the stimulation periods) using t-tests (alpha = 0.05). Each animal served as their own control.

## Results

We considered animals that completed all 4 nerve stimulation trials (n = 10) as experimental units and used them for full data analysis. Three animals were used to perform supplementary nerve transection trials. Two animals had blood perfusion measured using different LSCI parameters to test for aliasing. The remaining two animals died prematurely during the second stimulation trial and were excluded from analysis. Estrous stage, stimulation order, experiment number, and weight had no impact on results. Detailed experimental demographics can be found in Table 1. Experimental data and MATLAB scripts used in the data analysis are available online [35].

**Table 1.** Summary of experiment parameters

| Animal ID | Weight (kg) | First Nerve Target | Estrous Stage | Intact Spinal Cord |  | Spinalized |  |
| --- | --- | --- | --- | --- | --- | --- | --- |
| | | | | Pudendal Nerve MT (μA) | Tibial Nerve MT (μA) | Pudendal Nerve MT (μA) | Tibial Nerve MT (μA) |
| A | 0.29 | Tibial | Diestrus | 300 | 400 | 500 | 1000 |
| B | 0.29 | Tibial | Inconclusive | 3000 | 3600 | 2100 | 1800 |
| C | 0.29 | Tibial | Proestrus | 500 | 300 | 500 | 300 |
| F | 0.29 | Pudendal+ | Metestrus | 500 | 1300 | 500 | 3400 |
| G | 0.30 | Tibial+ | Metestrus | 400 | 250 | 400 | 900 |
| J | 0.28 | Pudendal | Proestrus | 300 | 600 | 500 | 5000 |
| K | 0.26 | Pudendal | Proestrus | 200 | 400 | 1000 | 400 |
| M | 0.25 | Tibial | Inconclusive | 300 | 750 | 600 | 600 |
| N | 0.26 | Pudendal | Proestrus | 130 | 400 | 300 | 400 |
| O | 0.26 | Pudendal | Inconclusive | 750 | 1700 | 750 | 1500 |
| P | 0.26 | Tibial | Estrus | 140 | 250 | 170 | 700 |
| Q | 0.26 | Pudendal* | Estrus | 170 | - | - | - |
| T | 0.25 | Tibial | Diestrus | 260 | 110 | 400 | 530 |
| U | 0.26 | Pudendal* | Metestrus | 110 | - | - | - |
| V | 0.26 | Pudendal* | Inconclusive | 600 | - | - | - |

MT: motor threshold, \*: Nerve Transection Experiment, +: Sampling Frequency Experiment

### Intact Spinal Cord Recordings

During pudendal nerve stimulation trials (Figure 2a), the average VBP at baseline was 184 ± 44 APU, which increased to 331 ± 129 APU during stimulation (+80.9 ± 64.2% change from baseline) and fell to 185 ± 55 APU after stimulation. Similarly, the average ABP was 172 ± 40 prior to stimulation, 348 ± 164 during stimulation (+97.6 ± 74.4%), and 169 ± 44 APU after stimulation ended. The average ITBP was 94 ± 33, 154 ± 67 (+67.0 ± 63.7%), and 92 ± 35 before, during, and after pudendal stimulation. The average blood perfusion during pudendal nerve stimulation for all ROIs was significantly higher than at baseline and after stimulation (p < 0.01).

**Figure 2.**
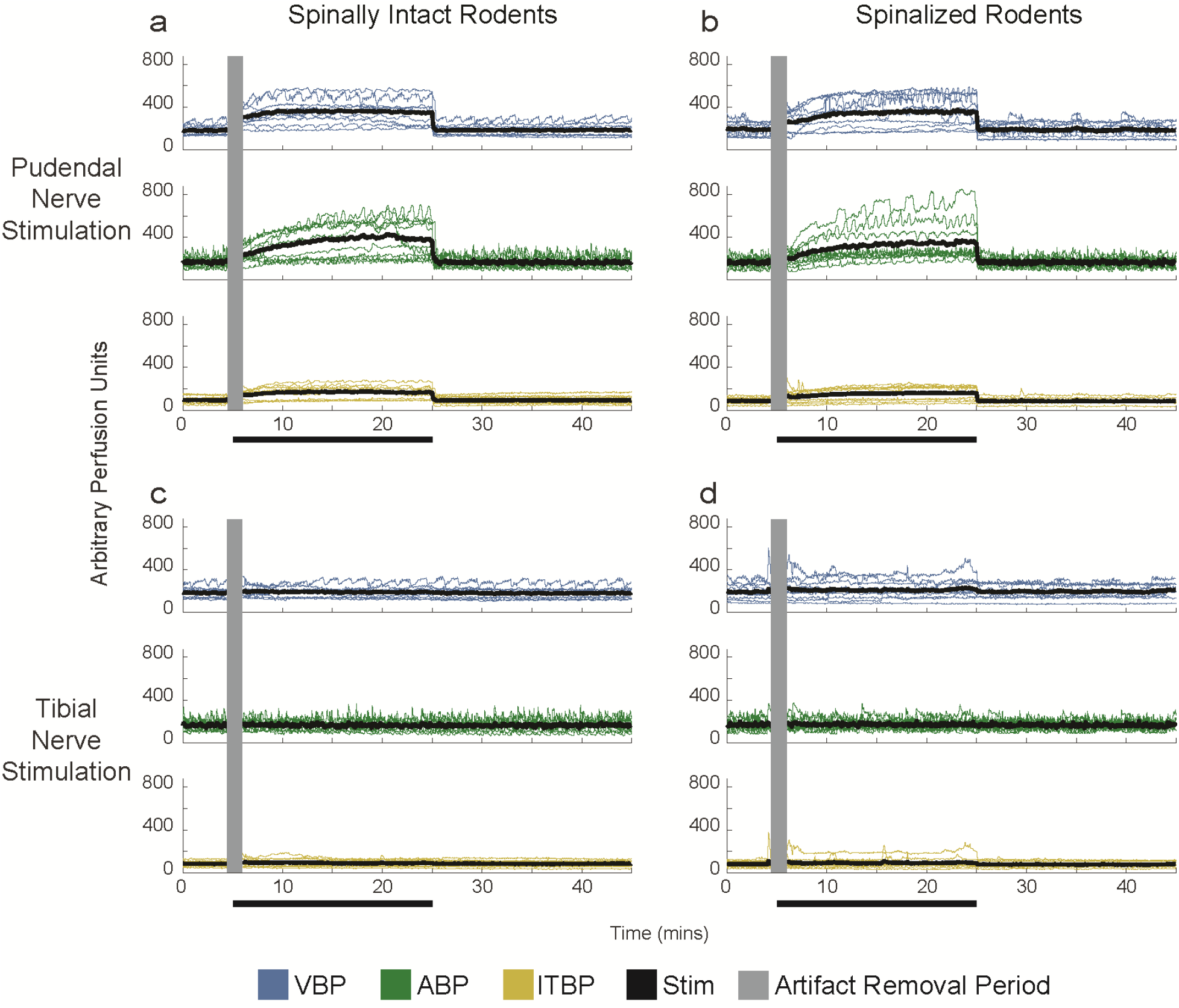
Individual (thin, colored lines) and average (solid black line) blood perfusion responses for all animals that completed the entire nerve stimulation protocol (n=10). a) Pre-spinalization pudendal stimulation trials. Blood perfusion returned to baseline after stimulation was turned off at 25 minutes. b) Pudendal stimulation trials after spinalization follow similar trends as pre-spinalization trials. Tibial nerve stimulation trials for intact (c) and spinalized (d) trials.

The average VBP for tibial nerve stimulation trials (Figure 2c) was 183 ± 50 APU prior to stimulation, 192 ± 48 APU during stimulation (+5.7 ± 11.9%), and 181 ± 52 APU after stimulation ended (Figure 3c). For the same trials, ABP was 163 ± 42, 163 ± 42 (+0.4 ± 8.0%), and 162 ± 47 APU before, during, and after stimulation. ITBP had mean values for the three periods of 85 ± 26, 93 ± 34 (+7.8 ± 22.5%), and 85 ± 31 APU. Tibial nerve stimulation did not elicit significant changes in any ROIs during stimulation, and the average blood perfusion was significantly lower in all ROIs compared to pudendal nerve stimulation (p < 0.001). Neither pudendal nor tibial stimulation elicited changes in perineal blood perfusion that persisted after stimulation was turned off.

**Figure 3.**
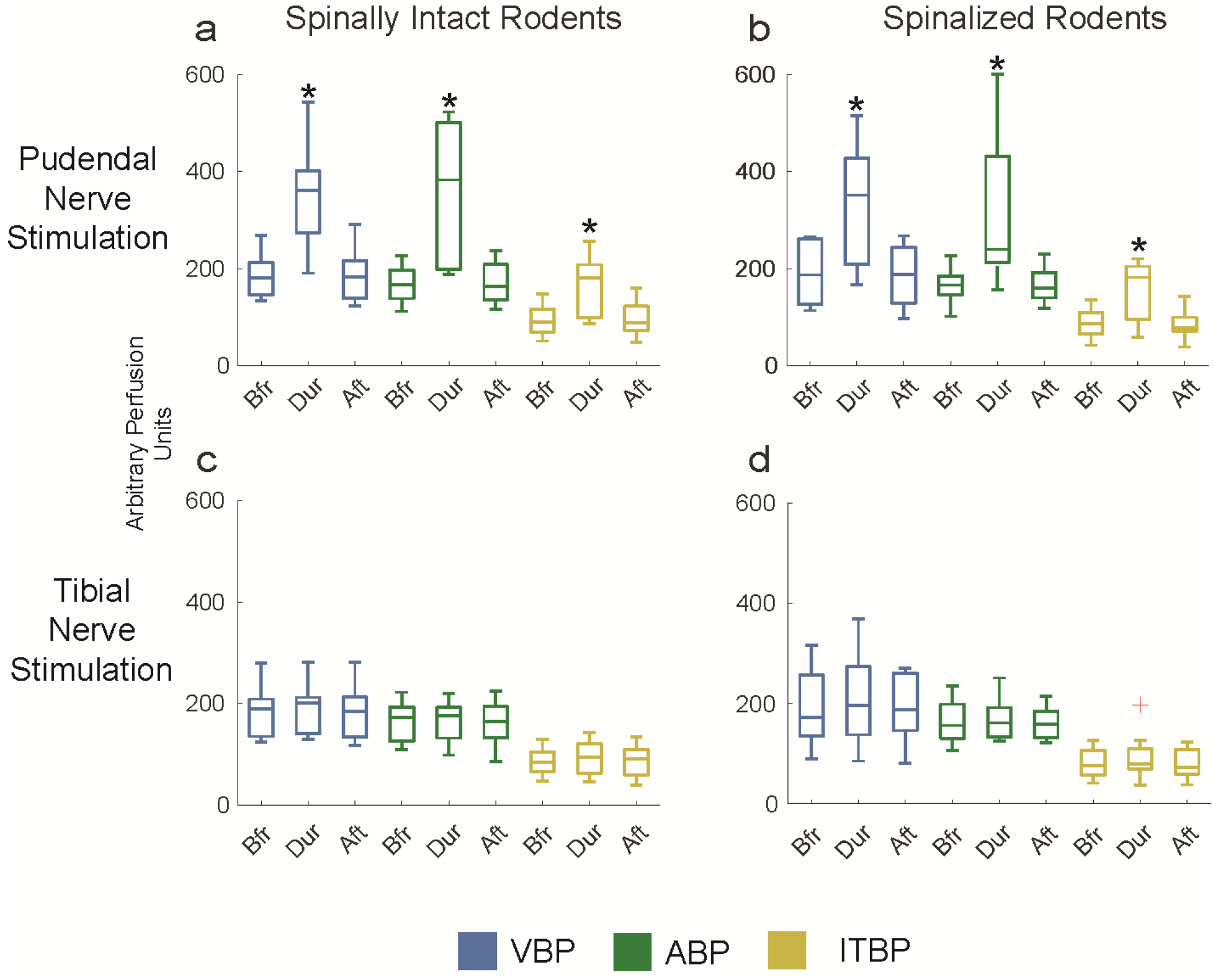
Boxplots of blood perfusion units across trial epochs (before, during, and after stimulation). Boxplot central lines give the median, edges indicate the interquartile ranges, whiskers represent range, and the cross denotes an outlier. Pudendal nerve stimulation trials (a, b) found elevated blood perfusion across all three ROIs during stimulation. There were no changes during tibial stimulation trials (c, d). * denotes significant difference (p < 0.05) between before and during stimulation epochs for an ROI.

### Post-Spinalization Recordings

Pudendal nerve stimulation trials after spinalization (Figure 2b) had an average VBP of 193 ± 61, 337 ± 131 (+74.5 ± 42.8% change from post-spinalization baseline), and 188 ± 65 APU before, during and after stimulation respectively. ABP for the same trials was 165 ± 39, 316 ± 161 (+86.6 ± 59.1%), and 167 ± 37 APU. Average ITBP was 88 ± 30, 153 ± 63 (+74.4 ± 60.7%), and 86 ± 28 APU (Figure 3b). There were no significant differences between average blood perfusion before and after spinalization for each ROI.

Tibial nerve stimulation trials had no significant differences after spinalization (Figure 2d). VBP was 193 ± 76, 211 ± 87 (+9.1 ± 11.8%), and 196 ± 65 APU prior to, during, and after stimulation. ABP was 168 ± 46, 171 ± 41 (+2.6 ± 7.8%), and 163 ± 32 APU for the same epochs. ITBP was 83 ± 29, 93 ± 45 (+10.0 ± 21.6%), and 79 ± 28 APU (Figure 3d).

### Sampling Frequency Experiments

In the sampling frequency experiments, we sampled blood flow using the LSCI system at 1 Hz for the first 4 minutes (no stimulation) and then at 25 Hz for 2 minutes (1 minute stimulation off, 1 minute on) to capture any artifact resulting from stimulation being turned on. We then reduced the sampling rate to 1 Hz for 18 minutes during stimulation. Then, we increased the sampling rate to 25 Hz for 3 minutes (2 minutes stimulation on, 1 minute off) to capture the transition during stimulation cessation. Finally, we used a sampling rate of 1 Hz for the final 19 minutes (stimulation off). In these two experiments, we observed regular, gradual oscillations in blood perfusion for each ROI that occurred at 1–3-minute intervals and were 100 to 300 APU in amplitude (e.g., Figure 4b). These oscillations were similar in width and amplitude as occasional rapid spontaneous increases in blood flow observed in 10 trials across 4 experiments in the main cohort (e.g., Figure 4a). This similarity suggests that the rapid changes during these trials were an artifact due to the low sampling rate. The stimulation artifact period during these experiments had the same features as in other experiments, with a brief, high signal period (generally 1 second or less in length), supporting our plan of removing that period during analyses.

**Figure 4.**
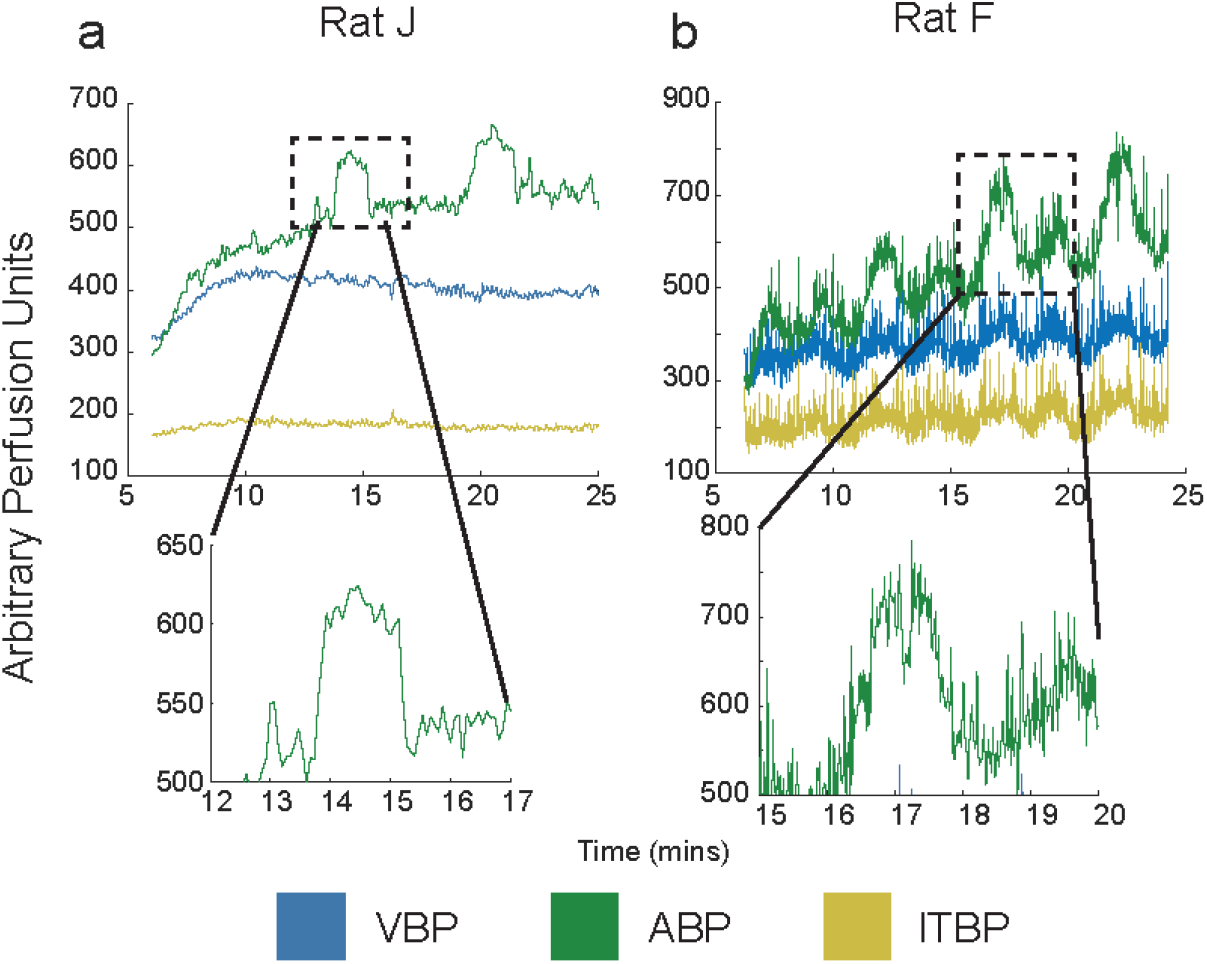
Intact spinal cord pudendal nerve stimulation trial periods of an experiment with abrupt increases in blood perfusion (4a) and an experiment with a higher LSCI sampling rate (4b). Top panels depict VBP, ABP, and ITBP waveforms during stimulation (0.25 and 1 Hz sampling frequency for Rat J (4a) and Rat F (4b) respectively). Bottom panels show zoomed-in plots of an individual signal oscillation in each trial of similar width (∼ 2 minutes) and amplitude (150-200 APU).

### Nerve Transection Experiments

We performed three experiments to examine the contributions of afferent and efferent pathways to the blood perfusion response to pudendal nerve stimulation. For the two pudendal nerve transection experiments with full cuts (one proximal and one distal to the stimulation location), the increases in VBP, ABP, and ITBP were abolished and there were no changes in blood perfusion for the duration of the trial. For the partial pudendal nerve transection experiment, there were still small, but noticeable increases in VBP, ABP, and ITBP during stimulation (+35.4%, +17.0%, and +43.8% change from baseline) which returned to baseline after stimulation was ceased. After complete pudendal nerve transection, we observed no changes in blood perfusion during stimulation for any of the ROIs (+0.9%, −6.5, and +1.0% change for VBP, ABP, and ITBP respectively).

## Discussion

In this study, we examined perineal blood flow changes in response to pudendal and tibial nerve stimulation. Additionally, we studied the impact of spinalization on these blood flow responses. This is the first study of its kind to use LSCI to measure perineal blood flow in animals. In intact animals, we saw a transient increase in perineal blood flow (VBP and ABP) during pudendal nerve stimulation that returned to baseline immediately after stimulation was turned off (Figure 3). Tibial nerve stimulation in intact animals did not impact blood flow during or after stimulation was delivered (Figure 2c). After spinal cord transection, pudendal and tibial nerve stimulation had the same effect as during intact spinal cord trials (Figure 2b/d).

Perineal blood perfusion increases during pudendal nerve stimulation are likely attributable to the direct activation of motor efferents, causing the anal and urethral sphincter to contract immediately after stimulation onset [36]. This muscle activation causes a generalized increase of blood flow to the entire perineal region. Spinal cord transection (Figure 3b) had no impact on the pudendal stimulation response, excluding the involvement of a supraspinal pathway. Transecting the pudendal nerve distally abolished the response immediately, indicating that the neural pathways involved are likely direct efferents, and not a genital somato-visceral spinal reflex. Furthermore, studies demonstrating a potential genital somato-visceral reflex that increases vaginal blood perfusion have a response that persists after stimulation is turned off [19, 20], which our VBP responses did not. Transecting the pudendal nerve proximally in one experiment unexpectedly abolished the perineal blood perfusion response. It is possible that we damaged the nerve during suture placement or transection. Experimental limitations prevented a repetition of this trial. Other experimental limitations include our use of anesthesia, which may have dampened the effect of stimulation on vulvar blood flow, and the duration of stimulation. It is possible that repeated or longer stimulation sessions are required to evoke a non-direct response, increasing vulvar blood perfusion.

Prior preclinical pudendal nerve stimulation studies have used short duration (20 seconds) and long duration (30 minutes) stimulation and measured the blood perfusion response using laser doppler flowmetry (LDF) probes placed on the interior vaginal wall [19, 20]. Both studies found an increase in vaginal blood flow that either began or was transiently maintained after stimulation was turned off. Vaginal blood flow gradually increased, peaked, and gradually returned to baseline during a 15-second to two-minute period. Preclinical experiments have also demonstrated the potential for tibial nerve stimulation to modulate vaginal blood flow [18, 37] and that estrogen is necessary to preserve this response [37]. It is not unexpected that a balanced gonadal hormone milleu is necessary for a genital blood flow response, however little research has been done to assess the direct impact of peripheral nerve stimulation on gonadal hormones. One such study found that tibial nerve stimulation did not have an impact on serum estradiol [37], and we hypothesize that pudendal nerve stimulation may not have a direct impact on gonadal hormones either. We expected to see increases in perineal blood perfusion in our experiments, however neither pudendal nor tibial nerve stimulation had an effect on perineal blood perfusion that persisted beyond the stimulation epoch. The vagina receives its blood supply from the uterine, vaginal, and internal pudendal arteries [17]. In contrast, the perineal region, which includes the vulva and anus, is only supplied by the internal pudendal arteries [17]. It is possible that pudendal nerve stimulation elicited changes in blood perfusion of internal tissues, such as the vagina, that could not be measured using LSCI.

Genital blood flow in women has been most commonly assessed with vaginal photoplethysmography (VPP) [25]. However, more researchers are using non-invasive techniques such as Laser Doppler Imaging (LDI) [26, 28] and LSCI [38] because they report higher levels of agreement between genital arousal, subjective arousal, and lubrication. Studies using LDI or LSCI to assess genital arousal have thus far been made in clinical settings in which participants receive visual sexual stimulation. Nerve stimulation may not be sufficient to evoke a lasting vulvar blood perfusion response in an anesthetized model of genital arousal. This suggests that vulvar blood perfusion needs further investigation before it is used in anesthetized animals as a direct measurement of genital arousal.

The absence of blood perfusion changes during tibial nerve stimulation (Figure 2, 3) is not entirely unexpected, as the tibial nerve innervates the lower leg and does not directly innervate the perineal region [39]. Spinal cord transection did not change the lack of response. Researchers have previously found that vaginal blood flow, measured with LDF, can increase in response to tibial nerve stimulation [18, 37]. The absence of blood perfusion changes in these experiments could be attributed to a difference in recording areas (vagina vs vulva), just as in pudendal nerve stimulation trials. Clinical studies have demonstrated that tibial nerve stimulation can improve bladder dysfunction in both non-neurogenic and neurogenic populations in weekly percutaneous stimulation sessions [15, 40]. In those studies, tibial nerve stimulation improved bladder storage, a function controlled by the sympathetic nervous system. However, healthy sexual functioning requires coordination of the sympathetic, parasympathetic, and somatic nervous systems and it is the parasympathetic pathways that cause smooth muscle relaxation and vaginal vasocongestion. Different stimulation parameters or repeated stimulation sessions (as is common for clinical treatment) may cause a change in VBP similar to perfusion changes observed in vaginal tissue.

We did not expect to see changes in ITBP during pudendal nerve stimulation, as the pudendal nerve does not directly innervate the superficial regions of the inner thigh. Spinalization did not have an impact on the average ITBP, across all stimulation epochs (Figure 3). Distal pudendal nerve transection abolished the increase in ITBP observed in intact animals during pudendal nerve stimulation. The pudendal nerve does not directly innervate any regions of the leg, but one cadaver study discovered that the sciatic artery can arise from the internal pudendal artery [41]. It is possible that activation of motor efferents caused blood flow to increase in pudendal arteries that then branch off and supply the inner thigh. Thus, the increases in ITBP are likely attributable to general blood flow to the region due to the contraction of the pelvic floor muscles placing an increased demand on blood supply, further supporting our conclusion that the increases in vulvar blood perfusion are due to somatic activation of the pelvic floor musculature.

This study sought to assess the impact of two potential treatment modalities for FSD (pudendal and tibial nerve stimulation) on perineal blood perfusion. We theorize that improvements in FSD symptoms are related to improvements in genital arousal and that it is possible to modulate genital arousal via pudendal or tibial nerve stimulation. Both treatment modalities have demonstrated potential in treating FSD. The dorsal genital nerve, a distal branching of the pudendal nerve, is a promising target because it can be accessed superficially, and stimulation has shown to modulate sexual function [16]. Tibial nerve stimulation is ideal for similar reasons as it is both superficial and easy to access, but more work is necessary to understand how it modulates genital blood flow. However, this preliminary study did not find any lasting changes in genital arousal as measured by perineal blood perfusion. We suspect that this contrast with prior preclinical studies is due to a difference in blood perfusion location and that the responses we did observe were due to muscle contractions requiring an increased blood supply to the region. An improved animal model for evaluating genital blood perfusion would incorporate simultaneous measurements of vaginal and vulvar blood perfusion. This model is necessary to understand the complete hemodynamics of different genital structures, in particular the relationship between vaginal and vulvar blood perfusion during arousal.

## Acknowledgements

We thank Tim Brown of Moor Instruments for assistance with data collection and analysis, Eric Kennedy and Ahmad Jiman for their help in experiment preparation, and the University of Michigan Unit for Laboratory Animal Medicine for animal husbandry. This study was supported in part by National Institutes of Health Award T32NS115724.

## References

1. Kashdan TB, Goodman FR, Stiksma M, Milius CR, McKnight PE. Sexuality leads to boosts in mood and meaning in life with no evidence for the reverse direction: A daily diary investigation. Emotion. 2018;18(4):563–76.

2. Laumann EO, Paik A, Rosen RC. Sexual dysfunction in the United States: Prevalence and predictors. J Am Med Assoc. 1999;281(6):537–44.

3. Meston CM, Stanton AM. Understanding sexual arousal and subjective–genital arousal desynchrony in women. Nat Rev Urol. 2019;16:107–20.

4. Katz M, Derogatis LR, Ackerman R, Hedges P, Lesko L, Garcia M, et al. Efficacy of flibanserin in women with hypoactive sexual desire disorder: Results from the BEGONIA trial. J Sex Med. 2013;10(7):1807–15.

5. Clayton AH, Althof SE, Kingsberg S, Derogatis LR, Kroll R, Goldstein I, et al. Bremelanotide for female sexual dysfunctions in premenopausal women: A randomized, placebo-controlled dose-finding trial. Womens Health. 2016;12(3):325–37.

6. Kingsberg SA, Clayton AH, Portman D, Williams LA, Krop J, Jordan R, et al. Bremelanotide for the Treatment of Hypoactive Sexual Desire Disorder: Two Randomized Phase 3 Trials. Obstet Gynecol. 2019;134(5):899–908.

7. Jaspers L, Feys F, Bramer WM, Franco OH, Leusink P, Laan ETM. Efficacy and safety of flibanserin for the treatment of hypoactive sexual desire disorder in women: A systematic review and meta-analysis. JAMA Intern Med. 2016;176(4):453–62.

8. Cavalcanti AL, Bagnoli VR, Fonseca ÂM Pastore RA, Cardoso EB, Paixão JS, et al. Effect of sildenafil on clitoral blood flow and sexual response in postmenopausal women with orgasmic dysfunction. Int J Gynecol Obstet. 2008;102(2):115–9.

9. Laan E, Van Lunsen RHW, Everaerd W, Riley A, Scott E, Boolell M. The enhancement of vaginal vasocongestion by sildenafil in healthy premenopausal women. J Womens Health Gend Based Med. 2002;11(4):357–65.

10. Basson R, McInnes R, Smith MD, Hodgson G, Koppiker N. Efficacy and Safety of Sildenafil Citrate in Women with Sexual Dysfunction Associated with Female Sexual Arousal Disorder. J Womens Health Gend Based Med. 2002;11(4).

11. Siegel S, Noblett K, Mangel J, Bennett J, Griebling TL, Sutherland SE, et al. Five-Year Followup Results of a Prospective, Multicenter Study of Patients with Overactive Bladder Treated with Sacral Neuromodulation. J Urol. 2018;199(1):229–36.

12. Hull T, Giese C, Wexner SD, Mellgren A, Devroede G, Madoff RD, et al. Long-term durability of sacral nerve stimulation therapy for chronic fecal incontinence. Dis Colon Rectum. 2013;56(2):234– 45.

13. Yih JM, Killinger KA, Boura JA, Peters KM. Changes in Sexual Functioning in Women after Neuromodulation. J Sex Med. 2013;10:2477–83.

14. Pauls RN, Marinkovic SP, Silva WA, Rooney CM, Kleeman SD, Karram MM. Effects of sacral neuromodulation on female sexual function. Int Urogynecology J. 2007;18(4):391–5.

15. Van Balken MR, Verguns H, Bemelmans BLH. Sexual functioning in patients with lower urinary tract dysfunction improves after percutaneous tibial nerve stimulation. Int J Impot Res. 2006;18(5):470–5.

16. Zimmerman LL, Gupta P, O’Gara F, Langhals NB, Berger MB, Bruns TM. Transcutaneous Electrical Nerve Stimulation to Improve Female Sexual Dysfunction Symptoms: A Pilot Study. Neuromodulation. 2018;21(7):707–13.

17. Hannan JL, Cheung GL, Blaser MC, Pang JJ, Pang SC, Webb RC, et al. Characterization of the vasculature supply the genital tissues in female rats. J Sex Med. 2012;9(1):136–47.

18. Zimmerman LL, Rice IC, Berger MB, Bruns TM. Tibial Nerve Stimulation to Drive Genital Sexual Arousal in an Anesthetized Female Rat. J Sex Med. 2018;15(3):296–303.

19. Cai RS, Alexander MS, Marson L. Activation of Somatosensory Afferents Elicit Changes in Vaginal Blood Flow and the Urethrogenital Reflex Via Autonomic Efferents. J Urol. 2008;180(3):1167–72.

20. Rice IC, Zimmerman LL, Ross SE, Berger MB, Bruns TM. Time-Frequency Analysis of Increases in Vaginal Blood Perfusion Elicited by Long-Duration Pudendal Neuromodulation in Anesthetized Rats. Neuromodulation. 2017;20(8):807–15.

21. de Groat WC, Tai C. Impact of Bioelectronic Medicine on the Neural Regulation of Pelvic Visceral Function. Bioelectron Med. 2015;2(1):25–36.

22. Hoang Roberts L, Vollstedt A, Volin J, McCartney T, Peters KM. Initial experience using a novel nerve stimulator for the management of pudendal neuralgia. Neurourol Urodyn. 2021;40(6):1670–7.

23. Sipski ML, Alexander CJ, Rosen R. Sexual arousal and orgasm in women: Effects of spinal cord injury. Ann Neurol. 2001;49(1):35–44.

24. Anderson KD. Targeting recovery: Priorities of the spinal cord-injured population. J Neurotrauma. 2004;21(10):1371–83.

25. Woodard TL, Diamond MP. Physiologic measures of sexual function in women: a review. Fertil Steril. 2009;92(1):19–34.

26. Boyer SC, Bouchard KN, Pukall CF. Laser Doppler Imaging as a Measure of Female Sexual Arousal: Further Validation and Methodological Considerations. Biol Psychol. 2019;148:107741.

27. Jabs F, Brotto LA. Identifying the disruptions in the sexual response cycles of women with Sexual Interest/Arousal Disorder. Can J Hum Sex. 2018;27(2):123–32.

28. Styles SJ, MacLean AB, Reid WMN, Sultana SR. Laser Doppler perfusion imaging: A method for measuring female sexual response. BJOG Int J Obstet Gynaecol. 2006;113(5):599–601.

29. Giuliano F, Pfaus J, Balasubramanian S, Hedlund P, Hisasue S ichi, Marson L, et al. Experimental Models for the Study of Female and Male Sexual Function. J Sex Med. 2010;7(9):2970–95.

30. Marson L, Giamberardino MA, Costantini R, Czakanski P, Wesselmann U. Animal Models for the Study of Female Sexual Dysfunction. Sex Med Rev. 2013 Jul;1(2):108–22.

31. Boyer SC, Bouchard KN, Pukall CF. Laser Doppler Imaging as a Measure of Female Sexual Arousal: Further Validation and Methodological Considerations. Biol Psychol. 2019 Nov;148:107741.

32. Allers KA, Richards N, Sultana S, Sudworth M, Dawkins T, Hawcock AB, et al. I. Slow oscillations in vaginal blood flow: Alterations during sexual arousal in rodents and humans. J Sex Med. 2010;7(3):1074–87.

33. Kovacevic M, Yoo PB. Reflex neuromodulation of bladder function elicited by posterior tibial nerve stimulation in anesthetized rats. Am J Physiol - Ren Physiol. 2015;308(4):F320–9.

34. McKenna KE, Nadelhaft I. The organization of the pudendal nerve in the male and female rat. J Comp Neurol. 1986;248(4):532–49.

35. Bottorff EC, Bruns TM. Pudendal and tibial nerve modulated vulvar blood perfusion in anesthetized rats [Internet]. Open Science Framework. Available from: https://doi.org/10.17605/OSF.IO/YMDFV

36. Pacheco P, Martinez-Gomez M, Whipple B, Beyer C, Komisaruk BR. Somato-motor components of the pelvic and pudendal nerves of the female rat. Brain Res. 1989;490(1):85–94.

37. Xu JJ, Zimmerman LL, Soriano VH, Mentzelopoulos G, Kennedy E, Bottorff EC, et al. Tibial nerve stimulation increases vaginal blood perfusion and bone mineral density and yield load in ovariectomized rat menopause model. Int Urogynecology J. 2022; DOI: 10.1007/s00192-022-05125-5

38. Cyr MP, Pinard A, Dubois O, Morin M. Reliability of vulvar blood perfusion in women with provoked vestibulodynia using laser Doppler perfusion imaging and laser speckle imaging. Microvasc Res. 2019;121:1–6.

39. Badia J, Pascual-Font A, Vivó M, Udina E, Navarro X. Topographical distribution of motor fascicles in the sciatic-tibial nerve of the rat. Muscle Nerve. 2010;42(2):192–201.

40. Stampas A, Gustafson K, Korupolu R, Smith C, Zhu L, Li S. Bladder neuromodulation in acute spinal cord injury via transcutaneous tibial nerve stimulation: Cystometrogram and autonomic nervous system evidence from a randomized control pilot trial. Front Neurosci. 2019;13:119.

41. Georgakis E, Soames R. Arterial supply to the sciatic nerve in the gluteal region. Clin Anat. 2008;21(1):62–5.

